# Large-scale multi-omics analysis suggests specific roles for intragenic cohesin in transcriptional regulation

**DOI:** 10.1101/2021.09.29.462097

**Authors:** Jiankang Wang, Masashige Bando, Katsuhiko Shirahige, Ryuichiro Nakato

## Abstract

Cohesin, an essential protein complex for chromosome segregation, regulates transcription through a variety of mechanisms. It is not a trivial task to genome-widely assign the diverse cohesin functions. Moreover, the context-specific roles of cohesin-mediated interactions, especially on intragenic regions, have not been thoroughly investigated. Here we performed a comprehensive characterization of cohesin binding sites in several human cell types. We integrated epigenomic, transcriptomic and chromatin interaction data with and without transcriptional stimulation, to explore context-specific functions of intragenic cohesin related to gene activation. We identified a new subset of cohesin binding sites, decreased intragenic cohesin sites (DICs), which have a different function from previously known ones. The intron-enriched DICs were negatively correlated with transcriptional regulation: a subgroup of DICs were related to enhancer markers and paused RNA polymerase II, whereas others contributed to chromatin architecture. We implemented machine learning and successfully isolated DICs with distinct genomic features. We observed DICs in various cell types, including cells from cohesinopathy patients. These results suggest a previously unidentified function of cohesin at intragenic regions for transcription regulation.

## Introduction

Cohesin, a ring-shaped chromosome-bound protein complex, is required for holding sister chromatids together during certain phases of the cell cycle^1^. Recent studies suggest that cohesin also has a role in transcription regulation, maintenance of chromosome architecture^2^ and DNA repair^3^. The context-specific functions of cohesin have been investigated using chromatin immunoprecipitation followed by sequencing (ChIP-seq) and high-throughput chromosome conformation capture (Hi-C). The early study reported that most cohesin-binding sites overlap with CTCF to function as an insulator^4^. Conversely, a group of cohesin is reported to be CTCF independent and co-bind with tissue-specific transcription factors (TFs) to contribute to transcriptional regulation^5, 6^, possibly via the mediation of interactions between enhancers and promoters^7^. Other studies using Hi-C suggested that cohesin and CTCF are essential for forming topologically associated domains (TADs), the evolutionarily conserved chromatin domains that are hundreds of kilobases to several megabases in length^8, 9^. These studies focused on cohesin functions with respect to blocked interactions by insulation, or the formation of enhancer-promoter interactions which implicitly assume the positive regulation of gene expression. In contrast, a recent report showed that transcription elongation within gene bodies causes displacement of cohesin binding from chromatin, leading to disruption of cohesin-mediated chromatin loops^10^. Thus a subset of chromatin loops (either end of which may be located within the intragenic regions) mediated by cohesin is suggested to be negatively correlated with gene activation. Whereas modifications in intragenic regions affect transcription events^11, 12, 13^, the function of intragenic cohesin has hardly been discussed.

Mutations in cohesin complex and its loader (NIPBL) are observed in the cohesinopathy Cornelia de Lange syndrome (CdLS), a multisystem developmental disorder^14^, and in multiple types of cancers^15, 16^. Our previous study found that the diagnostic phenotype of CdLS is very similar to that of CHOPS syndrome^17^, which is caused by missense mutations in AFF4, a core component of the super elongation complex. Given the diverse functions of cohesin with respect to gene expression and chromatin folding, the underlying molecular mechanism responsible for the similarity between CdLS and CHOPS is yet unknown. Noteworthily, the CHOPS-related mutations in super elongation complex are also associated with transcriptional regulation by cohesin, indicating a common pathogenetic mechanism of cohesin in both CHOPS and CdLS. It can be a feasible hypothesis that intragenic cohesin has a distinct role that links the phenotypic similarities between CdLS and CHOPS.

Here, we conducted a large-scale epigenomic analysis to clarify the context-specific functions of cohesin sites, especially those in intragenic regions. To investigate the perturbation of cohesin binding sites by gene activation, we generated RNA sequencing (RNA-seq) and ChIP-seq data for cohesin and several TFs from MCF-7 cells with or without transcription stimulus. We also used many publicly available datasets including Hi-C, ChIP-seq, RNA-seq and chromatin interaction analysis by paired-end tag (ChIA-PET). First, we clarified that a subset of cohesin sites, which we refer to as ‘decreased intragenic cohesin sites’ (DICs), is distinct from the other groups of cohesin sites. Cohesin bindings on DICs are negatively correlated with transcriptional activation and locus compaction of chromatin. A part of DICs exhibit the high preference for enhancer marks and paused RNA polymerase II, whereas others contribute to chromatin architecture. Second, we performed ChIP-seq and RNA-seq with cohesin-depleted cells and confirmed that cohesin has an active function on DICs. Third, we applied machine learning and captured DICs with distinct epigenomic landscape, which is predictable across cell types. Finally, we conducted plenty of ChIP-seq in other cell types. Importantly, DICs can be observed across multiple cell types, including cells derived from patients with CdLS and CHOPS, in a cell-type-specific manner. The findings from our integrated analysis and machine learning approaches suggest an additional role for cohesin in the regulation of gene expression.

## Results

### Classification of DICs

MCF-7 cells, when treated with the transcriptional stimulator estradiol^18^, is a widely used model for investigating the transcription-dependent perturbation^5^. We prepared ChIP-seq data of cohesin (Rad21), cohesin loader (Mau2), CTCF and several TFs (ER, CBP, P300, AFF4, TAF1) from MCF-7 cells treated with vehicle (control, or Ctrl) or estradiol (E2, 45min). ChIP-seq and RNA-seq data generated in this study and their statistics are summarized in Supplementary Table 1. Other omics data used in this study are listed in Supplementary Tables 2-3. In total, we obtained 76,668 and 89,111 peaks as cohesin binding sites in the E2 and control data, respectively. Next, we examined the stimulation-dependent cohesin sites (Fig. 1a). Although the total number of cohesin peaks decreased after E2 stimulation, the proportion of peaks that increased (9.3%) after stimulation (log-fold change of peak intensity Mvalue^19^ > 0.5) was larger than the one that decreased (6.2%) (Mvalue < −0.5) (Fig. 1a, bottom). We also found that around 40% (36.3% for E2, 41.2% for control) of cohesin peaks did not overlap with CTCF peaks (Supplementary Fig. 1a). Such ‘cohesin-non-CTCF sites’ (hereafter, CNCs) overlapped with peaks of the enhancer markers P300 and CBP (Supplementary Fig. 1b), which is consistent with an earlier ChIP-seq study^5^. The cohesin loader Mau2 also preferred enhancer sites. In fact, 88.7% of CNCs with enhancer markers overlapped with Mau2, and Mau2 was localized at enhancer sites with and without cohesin binding (Supplementary Figs. 1c-d). This result implies the role of Mau2 for enhancer activity or for chromatin interaction, which can precede cohesin localization.

**Fig. 1.**
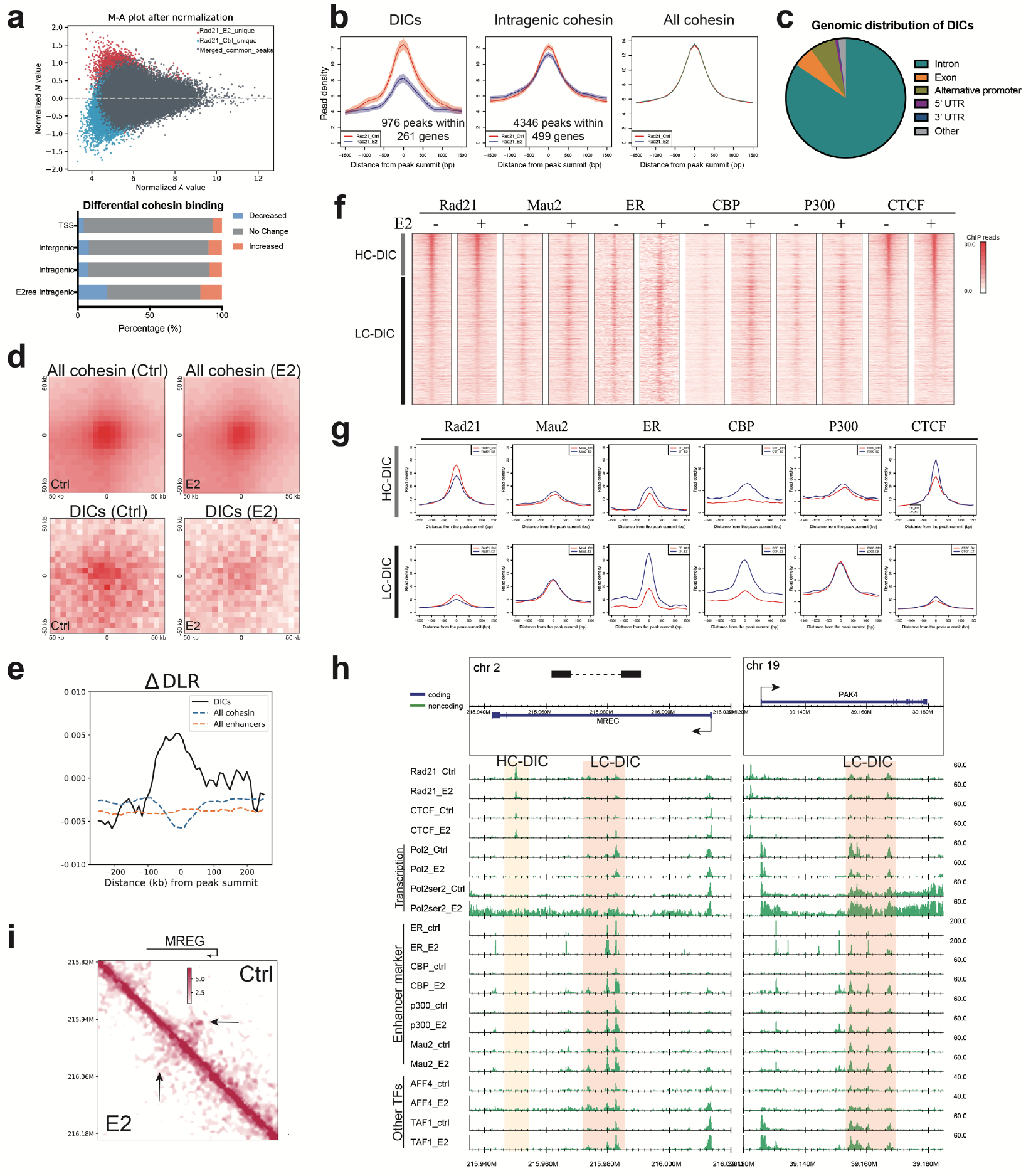
Classification of decreased intragenic cohesin (DIC) sites. **a**. Quantitative comparison of cohesin peak intensity between control (Ctrl) and estrogen (E2)-treated MCF-7 cells. Mvalue is the log_2_ (fold change) of normalized read densities under comparison. Avalue is the average signal strength of each peak. The lower panel shows the proportion of cohesin binding that decreased or increased. **b**. Average binding profiles on different cohesin sites. Red, control. Blue, E2 treatment. **c**. Genomic distribution of all DICs suggested most of DICs were located in introns. **d**. Aggregate peak analysis around DICs or all cohesin peaks at a 5-kb resolution. **e**. Chromatin compaction scores around DICs, all cohesin sites and all enhancers (summit ± 500 kb). ΔDLR (E2 vs. control) was calculated at a 25-kb resolution. **f**. Heatmap of ChIP-seq reads at DICs (peak summit ± 2.5 kb). CTCF (E2+) signal was used for sorting the order. DICs were divided into HC-DICs (gray bar) and LC-DICs (black bar). E2−, control; E2+, estrogen treated. **g**. Average binding profiles (summit ± 1.5 kb) of Rad21 and TFs on HC-DICs (upper) and LC-DICs (lower). Reads were normalized relative to the whole genome. The same *y*-axis scale was used for each protein. **h**. Read distribution of Rad21 and TFs around *MREG* and *PAK4* loci. Areas shaded in pink and light yellow indicate LC-DICs and HC-DICs, respectively. All reads were normalized relative to the whole genome. **i**. Hi-C contact map (5-kb resolution) around *MREG*. Arrows show the disappearance of chromatin interactions after E2 stimulation. The loop anchor is shown as a black bar in 1h.

We classified cohesin sites based on gene annotation information (Supplementary Fig. 1e). We defined ‘intragenic cohesin sites’ as those located within gene bodies, with the exception of transcription start sites (TSSs), transcription end sites (TESs) and alternative promoters. As a result, 13.8% of cohesin sites were identified as intragenic ones, 19.6% of which had enhancer activities. We did not observe a difference in the proportion of up- or down-regulated cohesin peaks between intergenic and intragenic sites (Fig. 1a, lower panel). To investigate the correlation of cohesin binding and transcription activation, we conducted the ChIP-seq of RNA polymerase II (Pol2), and RNA pol II CTD serine-2 phosphorylation (Pol2ser2) that represents transcription elongation activity^20^. We identified 499 E2-responsive genes for which the Pol2ser2 signal increased after E2 stimulation (Method). We then validated these genes by RNA-seq and confirmed that their expressions were mostly up-regulated by E2 stimulation (Supplementary Fig. 1f). Within the 499 E2-responsive genes, we identified 4,346 intragenic cohesin peaks, 976 (22.4%) of which were decreased after stimulation (Fig. 1b, Supplementary Fig. 1g). Because our main interest is the negative correlation between active transcription and the signal intensity of intragenic cohesin^10^, we focused on the intragenic cohesin sites for which the peak intensity decreased after E2 stimulation. Hereafter we refer to these sites as DICs. Among the E2-responsive genes, 53.5% (267/499) contained one or more DICs. We found that almost all (82.8%) DICs were located in intronic regions (Fig. 1c). This result implies a function of DICs on intron, whose mechanism remains unrecognized^21^.

Next, we investigated the correlation between decreased cohesin binding and the level of chromatin interaction using Hi-C data (GSE99451). Aggregate peak analysis (APA)^22^ showed that DICs-centered chromatin interactions were weakened by E2 treatment, whereas no difference was observed for all cohesin sites (Fig. 1d). These results suggested that at least some intragenic cohesin was required for chromatin loop formation, which were disrupted due to the induction of transcription according to previous study^10^. In contrast to the positive regulation of gene expression by CNCs^6^, DICs possibly function negatively with respect to gene expression. We then applied DLR (distal-to-local interaction ratio) and ICF (inter-chromosomal fraction of interactions) metrics^10^ to represent changes in locus-specific intra- and inter-chromosomal interactions, respectively. The differences (Δ) for DLR or ICF between two Hi-C samples represent chromatin compaction (negative value) or de-compaction (positive value). ΔDLR showed a positive value at DICs (Fig. 1e). In contrast, ΔDLR had a negative peak at all cohesin sites, whereas all enhancers showed no enrichment. Chromatin compaction at all cohesin sites could be explained by more frequent *cis*-regulatory interactions after estrogen stimulation. Conversely, DICs did not show a clear difference as compared with all cohesin sites for ΔICF (Supplementary Fig. 1h). These results suggested that DICs were involved in intra-chromosome decompaction that created a more open architecture around DICs.

### Classification of LC-DICs and HC-DICs

We next investigated the binding pattern of cohesin and other TFs, including the estrogen receptor (ER). We found that DICs could be clearly classified into two categories: HC-DICs (high CTCF binding) and LC-DICs (low CTCF binding), which co-localized with strong and weak CTCF peaks, respectively (Fig. 1f). LC-DICs had a higher probability of co-binding with many TFs as compared with HC-DICs (Figs. 1f-g). This tendency was similar, but not identical, to cohesin peaks in the other regions. For example, cohesin localized with strong CTCF on promoters, where many TFs also bound^6, 23^ (Supplementary Fig. 2a). A majority of intergenic cohesin sites (possibly insulator sites or TAD boundaries) did not show TF enrichment (Supplementary Fig. 2b). Moreover, the TFs on LC-DICs (Fig. 1g, Supplementary Fig. 2c, except Mau2 and P300), including 16 publicly available TFs (Supplementary Table 4), were increased after E2 treatment. This suggested that enhancer markers Mau2 and P300 localized to LC-DICs even before stimulation, whereas other TFs, including another enhancer marker CBP, were recruited by E2 stimulation. In addition, the weakened interactions were observed in both LC- and HC-DICs (Supplementary Fig. 2d).

Fig. 1h showed examples of two E2-responsive genes (*MREG* and *PAK4*; see Supplementary Fig. 2e for publicly available TFs). For instance, at the *MREG* locus, there were both HC-DICs, which co-localized with strong CTCF signals but with scarcely any TFs, and LC-DICs, which corresponded with the frequent binding of many TFs yet without strong CTCF signals. Overall, at LC-DICs, the peak intensity of cohesin decreased after E2 stimulation, whereas that of many TFs increased. Consistently, for the E2-activated gene *MREG* (Supplementary Fig. 3a), we could also clearly observe weakened interactions (Fig. 1i) and chromatin decompaction (Supplementary Fig. 3b).

More genomic characteristics were detected by motif analysis (Supplementary Fig. 3c). Not surprisingly, all types of cohesin showed the motifs of CTCF and CTCFL (BORIS). Specifically, LC-DICs were highly enriched for motifs of the forkhead box (FOX) protein family, which is responsible for remodeling chromatin structure^24^ and controlling transcription^25^. Of note, FOXA1 is a pioneer factor before ER activation in MCF-7 cells^26^. Meanwhile, HC-DICs showed motifs for transcription repressors including the tumor suppressor gene *HIC1*, implying a possible role for HC-DICs in transcription repression. Taken together, these results highlighted the unique features of LC-DICs and HC-DICs relative to other cohesin sites.

### Characterization of LC-DICs as enhancers

The bindings of the enhancer markers CBP and P300 were frequently observed at LC-DICs (Figs. 1f-h). We confirmed that a significantly higher percentage of LC-DICs overlapped with CBP binding as compared with other cohesin sites (Fig. 2a, Fisher’s exact test). In addition, LC-DICs were also enriched for enhancer markers H3K27ac and H3K4me1 as well as FANTOM5 enhancers^27^ (Fig. 2b, Supplementary Fig. 4a; publicly available data). In contrast, HC-DICs were not annotated as enhancers. Moreover, although intergenic cohesin also (including both CTCF-dependent and -independent) in conjunction with many TFs, they were not highly enriched for enhancer markers (Fig. 2a, Supplementary Fig. 4b). This is consistent with the finding that only 18% of intergenic cohesin was co-bound with CBP, which is reasonable because only a subset of intergenic cohesin sites serves as enhancers.

**Fig. 2.**
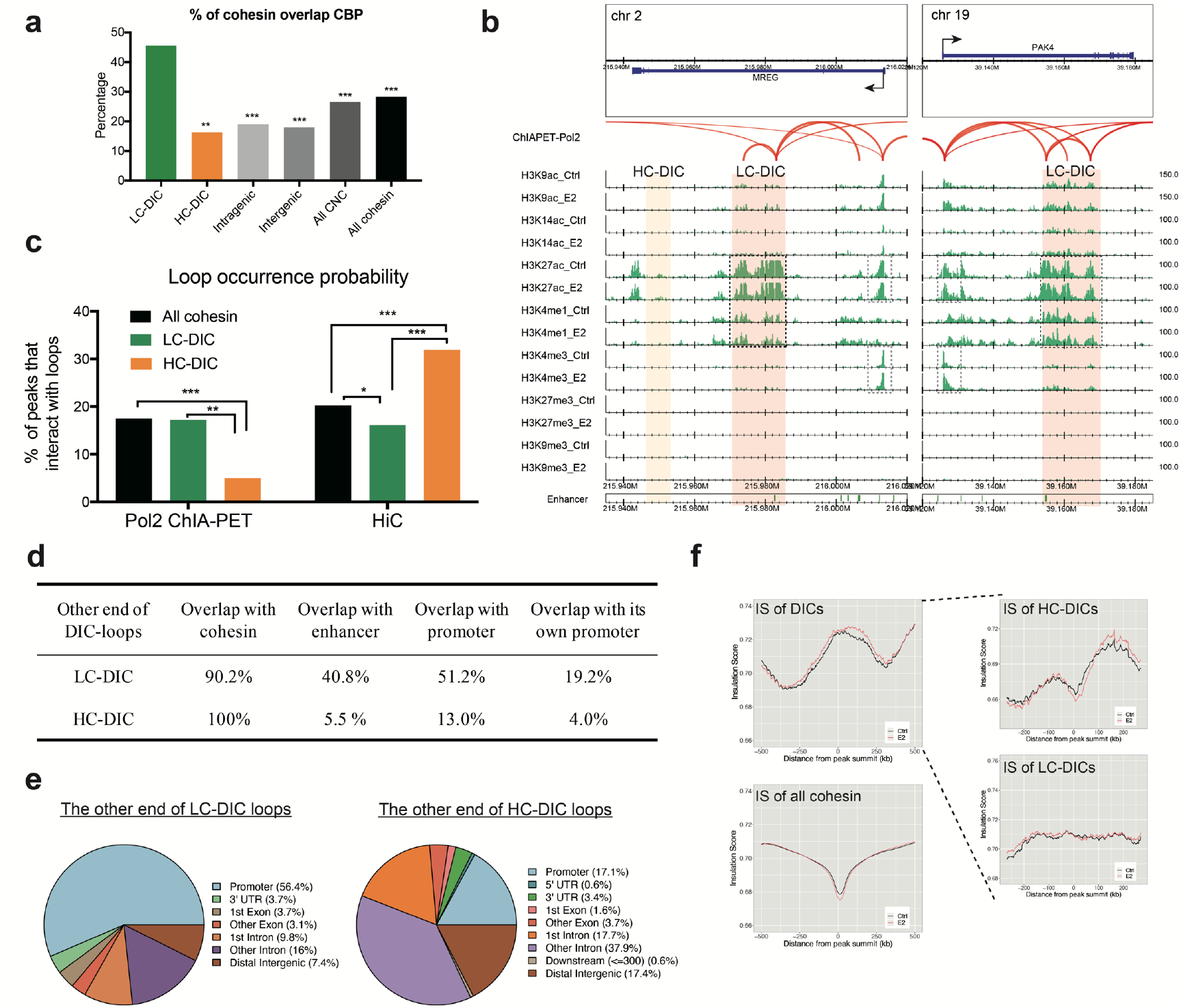
Enhancers and loops on DICs. **a**. The proportion of cohesin overlap with CBP peaks. Fisher’s exact test was used between LC-DICs and other cohesin sites. **p < 10^−5^, ***p < 10^−10^. **b**. Genomic binding of multiple histone markers. Enhancer markers H3K27ac and H3K4me1 (black dashed box) were enriched at LC-DICs. Gray dashed boxes indicate histone markers for promoters. Red arcs show Pol2-mediated loops. **c**. Loop occurrence probabilities on DICs, calculated based on Pol2 ChIA-PET or Hi-C data. We used Fisher’s exact test to do statistical comparisons. *p < 0.05, **p < 0.01, ***p < 0.001. **d**. The percentage of HiC loops (LC-DICs and HC-DICs) that interacted with cohesin, enhancers, promoters or the promoter of their host gene. **e**. Genomic distribution of the other end of DIC-loops as generated by ChIPseeker. **f**. Insulation scores (ISs) for each 25-kb genomic region around DICs or all cohesin sites (summit ± 500 kb). DICs were further divided (dashed line) into HC-DICs and LC-DICs.

### Characterization of chromatin loops on DICs

To explore DIC-mediated loops, we investigated what kind of chromatin loci interacted with LC-DICs and HC-DICs. Remarkably, when Pol2-mediated chromatin loops identified by ChIA-PET (GSE33664) were analyzed, the LC-DICs contained multiple Pol2 loops that interacted with the TSS of the host gene, whereas HC-DICs at the *MREG* locus did not have Pol2 loops (Fig. 2b, red arcs). To further investigate this tendency, we also analyzed DIC-anchored loops detected by Hi-C data (GSE99451). Interestingly, this result was in direct contrast to the Hi-C and ChIA-PET Pol2 loops (Fig. 2c, Fisher’s exact test). HC-DICs had a significantly lower occurrence probability with respect to Pol2-mediated chromatin loops as compared with LC-DICs and all cohesin sites. In contrast, HC-DICs exhibited a significantly higher occurrence with respect to Hi-C loops. We also compared loops with CTCF ChIA-PET data (GSE39495) and found that over 81% of HC-DICs overlapped with CTCF loops (27% for LC-DICs). This result suggested that LC-DICs were anchored by chromatin loops with Pol2 and other TFs to function as an enhancer in a CTCF-independent manner, whereas HC-DICs were more likely to interact with CTCF to form chromatin loops to participate in chromatin architecture independently of the Pol2 machinery.

We then investigated the other anchor sites of the DIC-mediated loops. The other anchor sites of DIC-mediated Hi-C loops also overlapped with cohesin, which also showed decreasing tendency (Supplementary Fig. 5a). As shown in Figs 2d-e, LC-DIC loops (ChIA-PET and Hi-C) mainly interacted with enhancers (40.8%) or promoters (51.2%), which were confirmed by highly enriched active histone markers (Supplementary Fig. 5b). We also observed that only a subset of LC-DIC loops (19.2%) interacted with the promoter of their host genes, suggesting that LC-DICs also contribute to the regulation of distant non-host genes, possibly as intragenic enhancer sites. In contrast, most of the HC-DIC loops did not interact with promoter or enhancer sites (Figs. 2d-e). Instead, over half of these sites interacted with intronic regions (Fig. 2e; example loci are shown in Supplementary Fig. 5c). In summary, these results suggested that LC-DICs participated in transcriptional regulation, whereas HC-DICs were more likely to connect the intronic regions of two genes.

We also examined the insulation score (IS) from Hi-C data, for which a lower value indicates more insulated regions, e.g., TAD boundaries. Although the IS profile showed clear valley at all, intergenic and intragenic cohesin sites (Fig. 2f, Supplementary Fig. 5d), it peaked at DICs (Fig. 2f, top left). Interestingly, the IS profile for HC-DICs showed bimodal peaks around small valley, whereas there was neither a peak nor valley at LC-DICs (Fig. 2f, lower right). These results suggested that LC-DICs possibly act as enhancers within TADs and that HC-DICs participate in the formation of boundaries, which is consistent with our loop analysis described above.

### Assessment of Pol2 stalling on DICs

Pol2 is released from promoter-proximal pausing to transcribe the entire gene body, although it may be temporarily paused by roadblocks within gene bodies^12^. To test whether DICs can function as roadblocks, we investigated the Pol2 enrichment at DICs using our Pol2 and Pol2ser2 ChIP-seq data (Supplementary Fig. 6a). Our Pol2ser2 and public global nuclear run-on sequencing data (GRO-seq, GSE99508) showed that transcription elongation was activated by E2 (Fig. 3a). Moreover, we found that Pol2 peaked at LC-DICs, and its intensity decreased after E2 stimulation (Fig. 3a-b), possibly due to the release of paused Pol2. This tendency toward a decrease in Pol2 binding was statistically significant as compared with the other cohesin sites (Fig. 3c). Importantly, ChIP-seq data treated with E2 for 30min, and public ChIP-seq datasets also showed the decreased Pol2 (Supplementary Figs. 6b-c). Given that bindings of most TFs were increased by E2 stimulation at LC-DICs (Figs. 1g-h), cohesin binding that decreased at LC-DICs was more likely to be accordant with Pol2, rather than TF bindings. In contrast, Pol2ser2 was increased on all DICs due to transcription activation (Fig. 3b, right panel). Pol2ser2 also exhibited peak-like enrichment at LC-DICs, which was increased by E2 stimulation (Figs. 3a-b). This is remarkable given that Pol2 enrichment at LC-DICs decreased significantly after E2 stimulation (Figs. 3b-c). In addition, whereas Pol2 binding on TSSs of DIC-host genes did not show any difference after E2 stimulation, the intensity of Pol2ser2 on TSSs increased (Supplementary Fig. 6d). These results support the model that Pol2 temporarily stalls within DICs, which function as roadblocks, and then is released by the loss of cohesin.

**Fig. 3.**
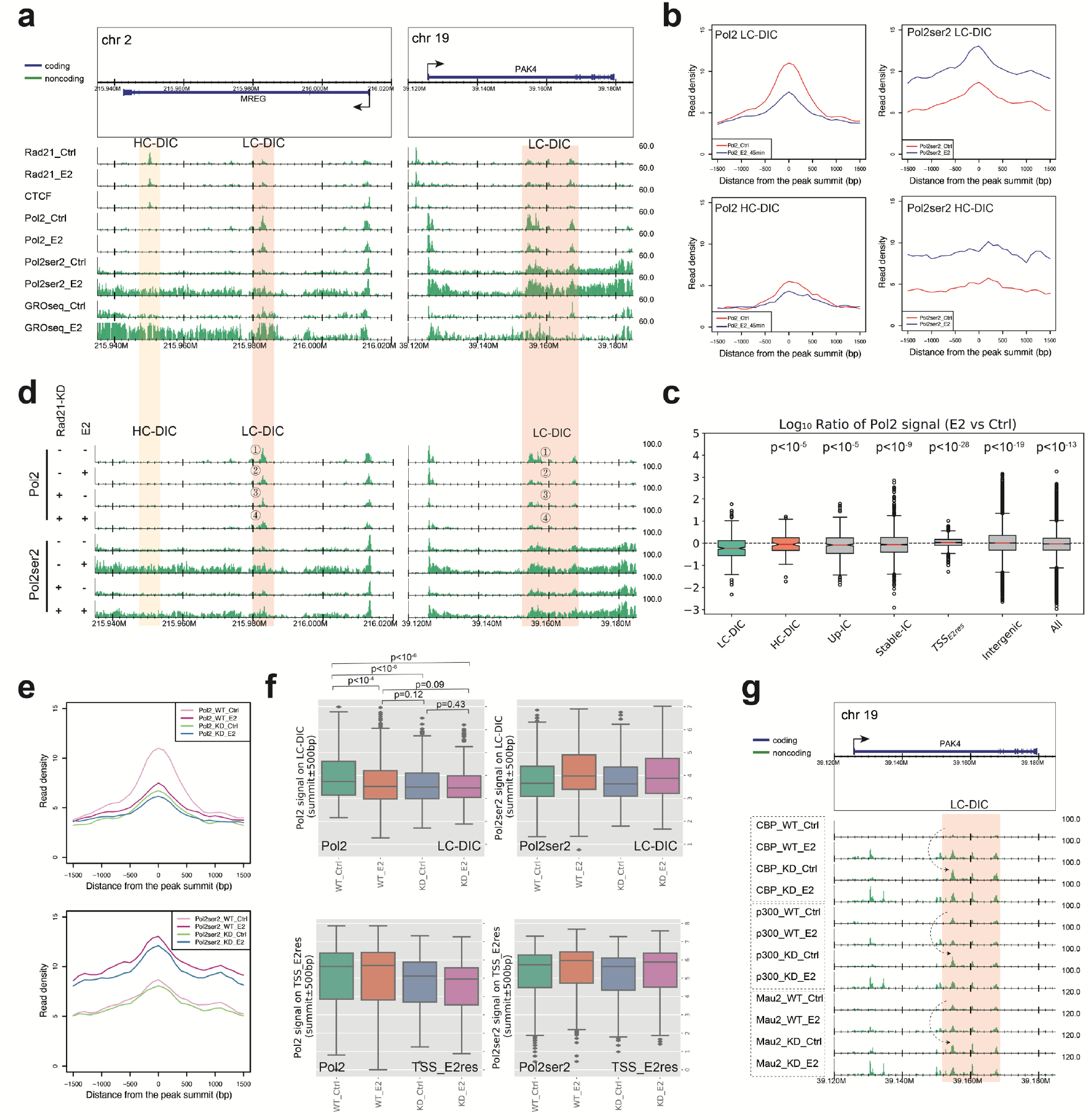
Pol2 pauses on LC-DICs. **a**. Genomic binding of Pol2 and Pol2ser2 and GRO-seq data for *MREG* and *PAK4* loci. **b**. Binding profiles of Pol2 and Pol2ser2 on LC-DICs or HC-DICs, under control and E2 conditions. Reads were normalized relative to the whole genome. **c**. The log_10_ value of the ratio E2/Ctrl with respect to the Pol2 signal on various cohesin sites; *p*-values (Mann–Whitney *U*-test) were calculated between LC-DICs and other cohesin sites. Up-IC, up-regulated intragenic cohesin; stable-IC, unchanged intragenic cohesin; TSS_E2res_, TSS of E2-responsive gene. **d**. Visualization of Pol2 and Pol2ser2 ChIP-seq data around *MREG* and *PAK4*. MCF-7 cells were treated as indicated. WT, wild type; KD, Rad21 knockdown; Ctrl, control; E2, estrogen. **e**. Pol2 and Pol2ser2 binding profiles around LC-DICs (summit ± 1.5 kb) under four different conditions. Reads were normalized relative to the whole genome. **f**. Quantitative comparison (Mann–Whitney *U*-test) of Pol2 and Pol2ser2 signals on LC-DICs or on TSSs of E2-responsive genes (TSS_E2res) under four different conditions. **g**. ChIP-seq results for CBP, P300 and Mau2 on *PAK4* gene in WT and Rad21 knockdown MCF-7 cells with the absence and presence of E2 stimulation. Dashed arrows show increased binding events in cohesin-deficient cells.

### Knockdown analysis of cohesin revealed that cohesin had a function at DICs

Although we observed cohesin binding at DICs that was synchronized with Pol2 binding and was negatively correlated with gene expression, there is still a possibility that cohesin is “passively” localized to DICs and therefore does not have any active role with respect to gene expression. To determine whether cohesin functions in the pausing of Pol2 at LC-DICs, we prepared Pol2 and Pol2ser2 ChIP-seq data in the absence and presence of Rad21-specific siRNA (siRad21) to generate Rad21 knockdown (KD; Supplementary Fig. 6e) and investigate the effect at DICs. Pol2 binding before E2 stimulation was significantly decreased by siRad21 (site 3 vs. 1 in Fig. 3d, KD_Ctrl as compared with WT_Ctrl in Figs. 3e-f) and had a similarly low level of binding in E2-treated wild-type cells (site 2 in Fig. 3d, WT_E2 in Figs. 3e-f). Thus, KD of cohesin indeed decreased Pol2 on LC-DICs, indicating that cohesin binding on LC-DICs is not passive but is required for Pol2 stalling. Pol2 binding in KD_Ctrl cells was not affected by E2-stimulation (site 4 vs. 3 in Fig. 3d, KD_E2 as compared with KD_Ctrl in Figs. 3e-f), possibly because Pol2 that was paused in WT_Ctrl cells had already been released in KD_Ctrl cells. Importantly, the effect of siRad21 on the Pol2 signal at TSSs of E2-responsive genes was distinct from LC-DICs, in which Pol2 binding did not change significantly after E2 stimulation in WT cells (from WT_Ctrl to WT_E2 in Figs. 3d and 3f) but decreased after siRad21 in stimulated cells (from WT_E2 to KD_E2 in Figs. 3d and 3f). These results also supported the model that the loss of cohesin binding causes the release of paused Pol2 on LC-DICs.

Interestingly, siRad21 did not largely affect Pol2ser2 binding. Pol2ser2 levels on LC-DICs showed no obvious difference between WT and siRad21 cells (Figs. 3d-f). In KD_Ctrl cells, there was no more stalling at LC-DICs, but there was also no stimulating effect of E2; thus Pol2ser2 did not change from WT_Ctrl to KD_Ctrl. In KD_E2 cells, transcription was activated by E2 stimulation but was limited by the loss of cohesin on TSSs, and thus Pol2ser2 binding changed slightly from WT_E2 to KD_E2. To explore changes in the expression level of genes that harbor LC-DICs after siRad21 treatment, we conducted RNA-seq with siRad21 (Supplementary Fig. 6f). Without E2 treatment, siRad21 did not significantly affect gene expression (*p*=0.23, KD_Ctrl as compared with WT_Ctrl). After E2 treatment, siRad21 moderately affected gene expression (*p*=0.005, KD_E2 as compared with WT_E2), possibly because only a subset of Pol2 that had paused on LC-DICs represented productive Pol2. We also quantitatively compared Pol2 and Pol2ser2 signals under four different conditions on various cohesin sites (Fig. 3f, Supplementary Fig. 6g). The results confirmed the significantly reduced bindings of Pol2 in WT_E2, KD_Ctrl and KD_E2 as compared with WT_Ctrl cells (Fig. 3f, Mann–Whitney *U*-test, one-sided). Such a tendency was distinct from those involving the TSSs of E2-responsive genes, up-regulated and non-changed intragenic cohesin, other cohesin sites, and also other enhancer sites. Our results suggested that the role of LC-DICs involves Pol2 pausing, which is different from the known roles of other cohesin sites.

In Figs. 1g-h, we showed the elevated binding of many TFs on DICs. To investigate whether the increased binding of multiple TFs is caused by a decrease in cohesin binding, we generated ChIP-seq data for CBP, P300 and Mau2 from siRad21 cells. Remarkably, a cohesin deficiency resulted in stronger binding of those TFs at LC-DICs, which surpassed the level in ER-stimulated WT cells (dashed arrow in Fig. 3g, Supplementary Fig. 7a-b). In contrast, there was little effect at the other intragenic enhancer site (Fig. 3g). Considering that E2 stimulation recruits TFs by estrogen responsive elements in WT cells, the increased binding of TFs in non-E2-stimulated siRad21 cells suggested that cohesin suppresses TF binding at LC-DICs in some way, and this suppression is removed by the loss of cohesin, which is followed by increased TF localization. Considering the chromatin de-compaction by E2 stimulation shown in Fig. 1e, the increased binding of TFs at LC-DICs can be, at least in part, explained by a more accessible chromatin structure near the LC-DICs, which is induced by the disruption of cohesin-mediated interactions.

### Machine learning analysis of DIC features

To confirm our findings, we implemented machine learning (ML) approaches (Supplementary Fig. 8a). We generated an integrated data matrix consisting of 175 features from genomic, transcriptomic and epigenomic data for all cohesin sites (Supplementary Table 5; Methods). Especially, this matrix includes features related to genomic location (e.g., intragenic or TSS) and to perturbation by E2 stimulation such as Mvalue and ΔDLR. Supplementary Fig. 8b showed a Pearson correlation heatmap followed by hierarchical clustering between all-by-all features for DICs or all cohesin sites. The 175 features resulted in clear clusters both among DICs and among all cohesin sites (dashed boxes of different colors). We annotated the clusters as promoter, enhancer, enhancer-promoter interaction (E-P), insulator and chromatin architecture. As compared with all cohesin sites, DICs showed lower co-binding tendency in the promoter cluster and higher co-binding in the enhancer and E-P clusters. This showed the effectiveness of our matrix in distinguishing DICs from other cohesin sites.

We then applied unsupervised k-means clustering (k = 10) to the matrix and obtained 10 clusters for all cohesin sites (cluster 0–9, Fig. 4a, Supplementary Fig. 10), among which only cluster 4 and cluster 7 showed intragenic cohesin binding that decreased after E2 stimulation, indicating the DIC-like clusters (Fig. 4b). We identified the following characteristics of cluster 4 (Fig. 4c, upper): 1) co-binding with tissue-specific TFs (e.g., ER and FOXA1), 2) enrichment of enhancer markers and Pol2, 3) relatively low intensity of cohesin and CTCF peaks and 4) chromatin de-compaction. Therefore, cluster 4 represented the LC-DIC-like cluster. In contrast, cluster 7 (Fig. 4c, lower) showed the following characteristics: 1) lack of TF co-binding, 2) high intensity of CTCF peaks and 3) highly related to topological boundaries and chromatin architecture features (e.g., TAD borders, Hi-C loops). Therefore, cluster 7 represented the HC-DIC-like cluster. Compared with “CNC-like” intragenic cohesin sites^5, 6^ (clusters 2, 3 and 8; Supplementary Fig. 10), cluster 4 (LC-DICs) co-localized only with enhancer markers and several master regulators (FOXA1, ER and GATA3), and therefore it is distinct from typical *cis*-regulatory modules (CRMs) at which many TFs co-localize. In contrast, cluster 7 (HC-DICs) consists of a cluster of intragenic cohesin sites that tend to be localized to open chromatin, are highly de-compacted and contain loops but are strongly negatively correlated with TFs. Therefore, they may be associated with a more universal chromatin structure that is required for proper gene transcription.

**Fig. 4.**
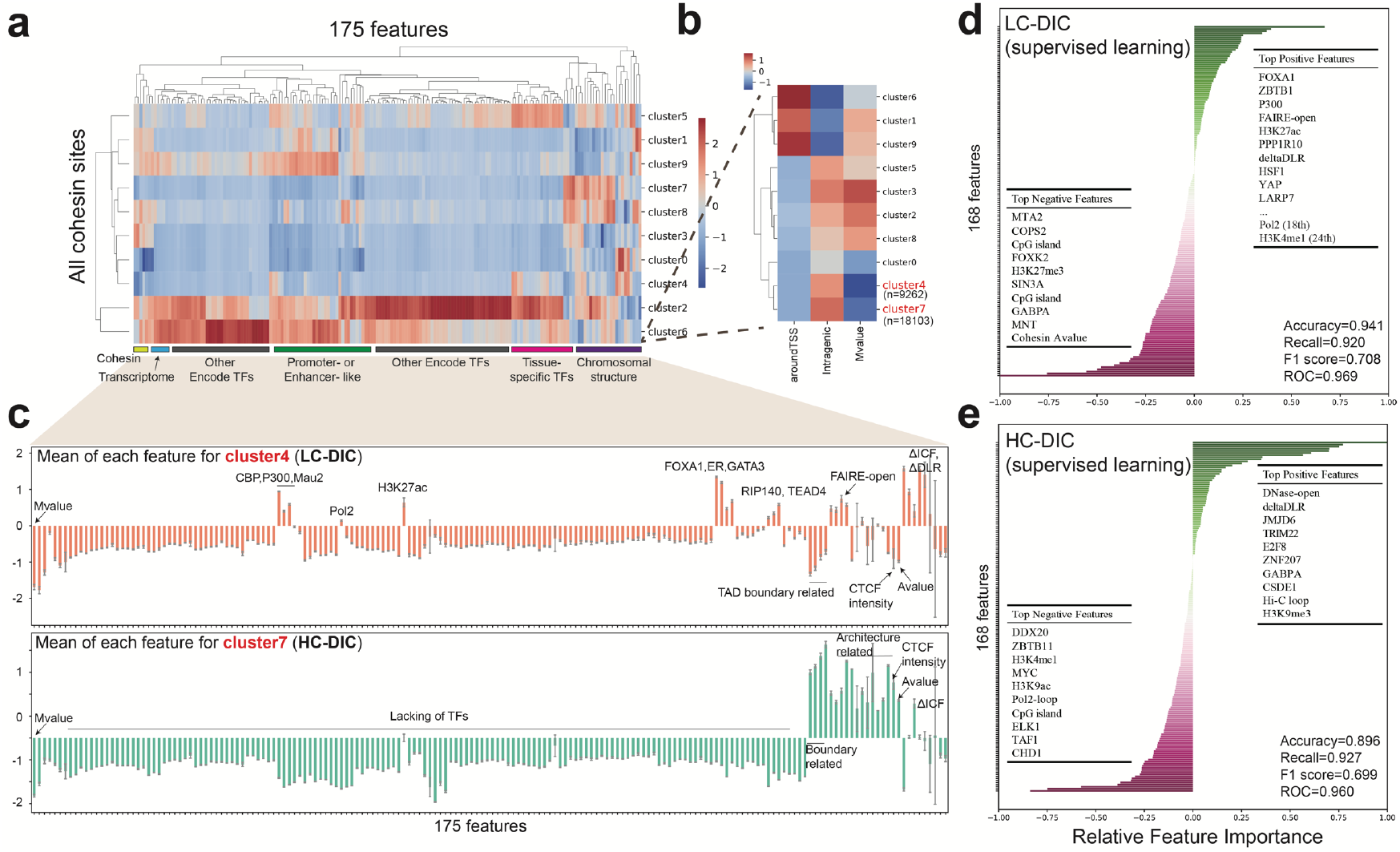
Machine learning methods to classify DICs. **a**. Cluster heatmap by k-means (k = 10) methods. The 175 features were roughly classified into different groups. **b**. Magnified view of the heatmap area shown in A indicates three important features for these 10 clusters: “aroundTSS” indicates whether a cohesin site was near TSS; “intragenic” indicates whether a cohesin site was within a gene body; “Mvalue” is the log_2_ ratio of Rad21 peak density between the E2 and control group. **c**. The mean of each feature for cluster 4 and 7. The *y* axis shows the normalized (z-score) values of binary and continuous features. Error bars represent 95% confidence intervals. **d-e**. The importance of individual features from the logistic regression model for (**d**) LC-DICs and (**e**) HC-DICs. Top positive and negative features are listed inside the plots. Model performance is indicated in the bottom right.

To further explore the characteristics of DICs, we applied modeling of supervised ML (logistic regression, support vector machine, and random forest) to predict LC- and HC-DICs from all cohesin sites in a binary manner (labeled by 0 or 1). In this analysis, the input matrix consisted of 168 features (features related to genomic location, cohesin changes and CTCF signal were excluded). We selected chromosomes 16 to 22 for testing, and the remaining chromosomes were divided into training and validation by five-fold cross-validation (see Methods). Because DICs are a small subset of all cohesin sites, we used SMOTE over-sampling^28^ to deal with such “imbalanced” classifications. The trained model that was based on logistic regression achieved the best performance overall as compared with the others (Supplementary Figs. 8c-d) and also performed adequately with the test data (Supplementary Fig. 8e). Finally, we identified important features for the prediction of LC-DICs and HC-DICs by calculating the relative feature importance from the trained model (Figs. 4d-e). LC-DICs were positively associated with 1) enhancer markers (H3K27ac, H3K4me1, P300); 2) Pol2 peaks, the Pol2-pausing regulator (LARP7)^29^ and a transcriptional repressor (ZBTB1)^30^; 3) tissue-specific regulators (FOXA1^26^, HSF1^31^, ER^18^). Both LC-DICs and HC-DICs were positively associated with open chromatin (FAIRE-open, DNase-open) and chromatin de-compaction (ΔDLR), and were negatively associated with H3K27me3 and CpG island levels. HC-DICs, in particular, showed positive features of Hi-C loops but negative features of Pol2 loops and TF binding, which is consistent with our analysis above, indicative of the TF-independent chromatin de-compaction. Taken together, the application of machine learning successfully isolated special subsets of cohesin sites (DICs) from all cohesin sites, which also provided us with additional characteristics by which they can be identified.

### Characterization of DIC tissue specificity

As DICs were enriched by many tissue-specific factors, we wondered whether our observations about DICs were consistent with other tissues or cell types. We generated Rad21 ChIP-seq for 293T cells (kidney), B-cells (lymphocytes), human skin fibroblast cells, RPE (retinal pigmented epithelium) cells, and HeLa cells (cervical cancer). Cohesin peaks at MCF-7 derived LC-DICs were more specific in MCF-7 cells, whereas cohesin peaks at HC-DICs were more ubiquitous across cell types (Figs. 5a-b). We then asked whether HC-DICs play a role in tissue-specific transcription. Across cell types, the peak intensities for Rad21 at HC-DICs were negatively correlated with transcription levels of their host genes (Fig. 5c, Supplementary Fig. 9a), suggesting that genes with stronger HC-DIC binding had lower transcription activities. Therefore, HC-DICs also showed the ability to impede transcription, which is consistent with our motif analysis above.

**Fig. 5.**
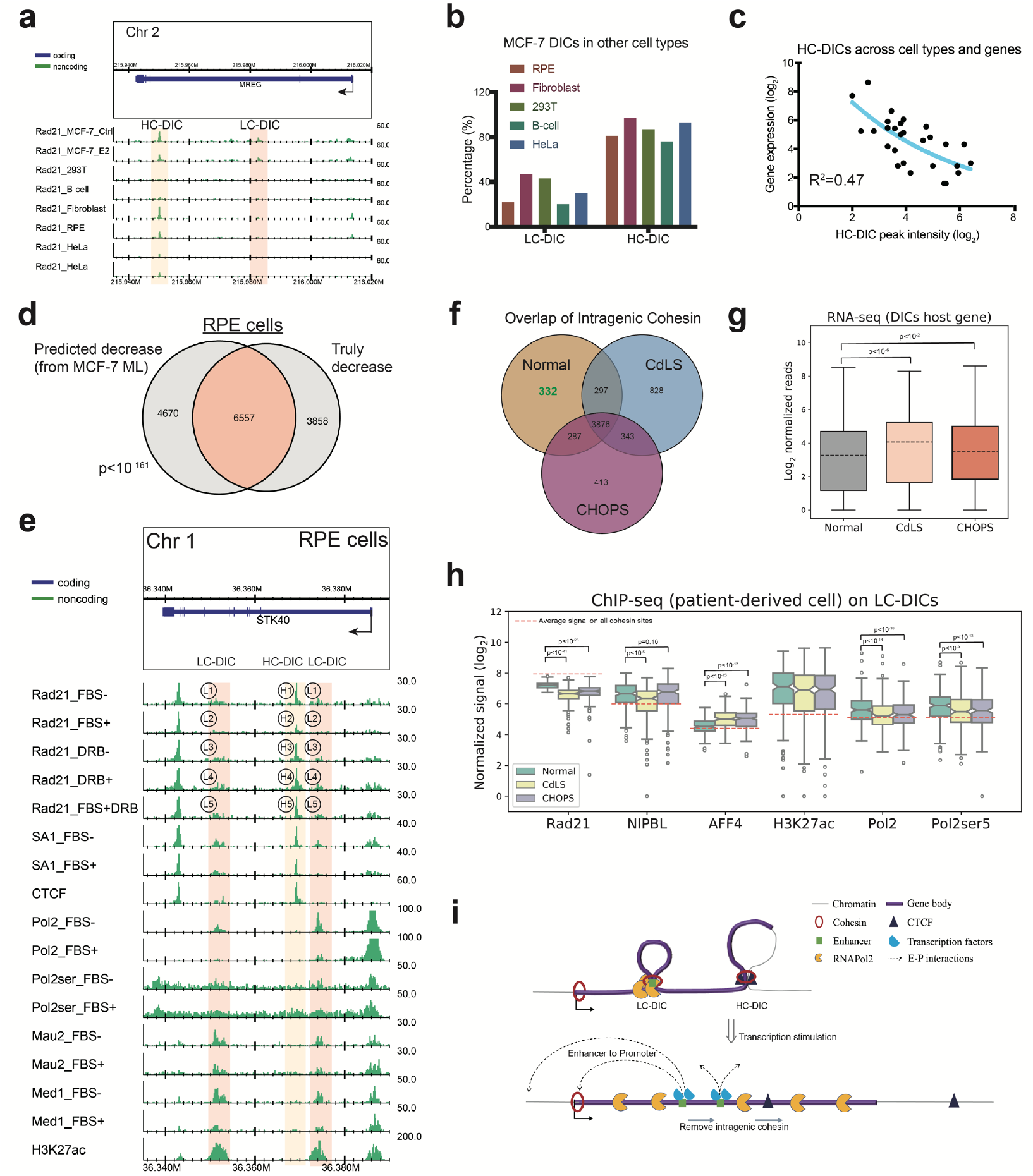
DICs in other cell types. **a**. Rad21 ChIP-seq of the *MREG* locus in various cell lines (MCF-7, 293T, B-cells, Fibroblasts, RPE, HeLa). HC-DICs were observed in other cells but not LC-DICs. **b**. The percentage of MCF-7 DICs that could be found in other cell types. **c**. Relationships between peak density of HC-DICs and the expression of their host genes. Gene expression data for the different cell types were obtained from the GTEx database. **d**. Overlap between DICs that were predicted relative to those that were experimentally isolated; *p*-values were calculated with the hypergeometric test. **e**. ChIP-seq data of the example locus in RPE cells. L1-5 indicated LC-DIC sites and H1-5 indicated HCDIC sites. **f**. Overlap of intragenic cohesin sites in normal and CdLS- and CHOPs-derived fibroblast cells. **g**. Expression of genes associated with DICs (paired t-test) in Normal, CdLS and CHOPs cells. **h**. Binding of Rad21, NIPBL, AFF4, H3K27ac, Pol2 and Pol2ser5 on LC-DICs. Dashed lines represent the average binding signal of each TF on all cohesin sites. Wilcoxon signed-rank test was used to compute *p*-values. **i**. Proposed model for our DICs. Briefly, intragenic loops formed by LC-DICs pause Pol2, whereas HC-DICs form CTCF-mediated loops. Transcription stimulation could displace DICs and disrupt DIC loops, consequently releasing the Pol2 pause. The exposed enhancer sites on LC-DICs then bind their respective TFs and interact with promoters.

To confirm whether DICs in other cell types also exhibit similar characteristics, we performed ChIP-seq experiments on RPE cells with FBS (fetal bovine serum) and DRB (5,6-dichloro-1-β-D-ribofuranosylbenzimidazole), which function as a stimulator and inhibitor of transcription^32^, respectively. First, we tested if the ML model trained by MCF-7 data was applicable to the RPE data. We used 25 features that were available for both MCF-7 and RPE cells to predict whether binding of an intragenic cohesin was decreased or not after transcription stimulation (Fig. 5d). The predicted DICs overlapped extensively with the experimentally determined ones (p < 10^−161^, hypergeometric test), indicating that DICs exhibited some common rules across cell types. Then we identified DICs of stimulation-responsive genes in RPE cells (Supplementary Fig. 9b). Similar to DICs in MCF-7 cells, RPE derived DICs also showed tissue-specific binding patterns (Supplementary Fig. 9c). FBS stimulation decreased the intensity of cohesin (Rad21 and SA1) at DICs (position H1 to H2, L1 to L2 in Fig. 5e; Supplementary Fig. 9d), whereas transcriptional inhibition by DRB increased it. In addition, further treatment with DRB (i.e., FBS+DRB) reverted the decrease in cohesin binding caused by FBS stimulation (H2 to H5, L2 to L5 in Fig. 5e; Supplementary Fig. 9d). Moreover, RNA Pol2 stalling and the release of paused Pol2 were also observed at LC-DICs in RPE cells (Fig. 5e, Supplementary Fig. 9e). In addition, LC-DICs, but not HC-DICs, co-bound with enhancer marks and several TFs. Thus, DICs are not a phenomenon associated with only breast cancer cells but also are found in normal human cell lines.

### Analysis of DICs in CdLS and CHOPS cells

Finally, we attempted to examine the participation of DICs in the observed phenotypes in individuals with CdLS and CHOPS. To this end, we generated ChIP-seq data from patient-derived fibroblast cells, together with the data that were generated in our previous study^17^. We overlapped the binding sites of intragenic cohesin in different cell types (Fig. 5f). Whereas most sites were shared among samples, there were 332 Rad21 sites that were absent in both CdLS and CHOPS cells, which we defined as DICs. RNA-seq analysis showed that host genes of DICs (both LC- and HC-DICs) in both CdLS and CHOPS cells showed a significant increase in transcription (Fig. 5g, Supplementary Fig. 9f; paired t-test), suggesting that the decreased cohesin binding at DICs was correlated with expression upregulation in both CdLS and CHOPS. We also assessed the binding of TFs at 185 LC-DICs and 147 HC-DICs (Fig. 5h, Supplementary Fig. 9g). Interestingly, at LC-DICs, the peak intensity of AFF4 (causative gene of CHOPS) increased in both CdLS and CHOPS cells, whereas lower binding of NIPBL (causative gene of CdLS) was observed only in CdLS cells (Fig. 5h). Enhancer marker H3K27ac was highly enriched but did not change among the three cell types, whereas Pol2 and Pol2ser5 (RNA pol II CTD phospho Ser5, which represents paused Pol2) decreased in both CdLS and CHOPS cells, which is consistent with our observations in MCF-7 and RPE cells. Taken together, this result suggests that DICs, especially LC-DICs, are involved in abnormal transcription associated with both CdLS and CHOPS. As both CdLS and CHOPs are involved in abnormal Pol2 regulation^33^, LC-DICs might offer a common pathogenetic mechanism. Based on the observations, we concluded that intragenic cohesin sites can be a good candidate to investigate and link the phenotypes of these two cohesinopathy disorders.

## Discussion

Cohesin is thought to be responsible for transcription regulation and chromatin folding. Several models have been proposed to explain its functions. Cohesin can mediate enhancer-promoter loops with the mediator complex or can function as a blocker between enhancer and promoter in conjunction with insulator factor CTCF^34^. Cohesin also participates in the formation of chromatin topological structures via the loop extrusion model^8^. A recent paper reported that transcription stimuli such as IFN-beta in THP-1 cells can displace cohesin from chromatin^10^, which attracted our interest. Here, we focused on intragenic cohesin, a subset of cohesin that was scarce, if not none, discussed in previous research. Of note, we emphasized the effect of negative regulation of gene expression by cohesin-mediated chromatin loops, whereas most of the previous studies implicitly assumed the positive regulation of expression by these loops. DICs were negatively associated with activated transcription and chromatin compaction. Whereas LC-DICs were highly enriched with enhancer markers and paused Pol2, HC-DICs were more involved in the features of chromatin architectures. Importantly, DICs could be found in multiple cell types, especially in CdLS and CHOPS cells, which partly supported the similarities between CdLS and CHOPS.

Chromatin interactions are required not only to facilitate transcription but also for Pol2 pausing^35^. By using siRad21 cells, we confirmed that the release of Pol2 was related to the loss of cohesin at LC-DICs, which supported our model that intragenic loops formed by cohesin paused Pol2 and that transcription elongation from TSSs could remove such cohesin and then release the paused Pol2 (Fig 5i). Notably, we did not observe significant changes in the expression of genes that host LC-DICs by siRad21 (Supplementary Fig. 6f). As most LC-DICs are located in intronic regions, Pol2 released from LC-DICs might be involved in accurate RNA splicing. Notably, a recent study suggested that intragenic enhancers, in addition to activating genes, also attenuate the transcription of their host genes during productive elongation^12^, which evokes the functional link between LC-DICs and Pol2 pausing. In contrast, HC-DICs showed a high preference for loop occurrence mediated by CTCF, possibly to play a role in topological boundaries (e.g., sub-TADs). Across different cell types and genes, the Rad21 signal at HC-DICs was negatively correlated with the expression of host genes, indicating the role of HC-DICs in restraining transcription. Whereas we observed that more than half of HC-DIC–mediated loops anchored intronic regions of two genes, it was difficult to infer the biological meaning of this because HC-DICs scarcely overlapped with any other TFs. Further biological approaches such as genome editing of HC-DICs of activated genes could be promising in the future.

Modifications at intragenic regions affect transcription events. For instance, intragenic methylation can prevent spurious transcription initiation^11^; Intragenic microRNAs affect the expression of their host genes^13^. Here we present the first study that focused on intragenic cohesin sites. We also used penalty linear regression followed by univariate linear regression to better understanding the changes of cohesin bindings in intragenic regions (see Method and Supplementary Fig. 8f-h). Apart from the decreased ones, the increased intragenic cohesin sites seemed to be also correlated with many important features, as they are positively predicted by ER and several TFs. Although we characterized intragenic cohesin sites that showed decreased bindings in this study, all types of intragenic cohesin might have a role in transcription regulation. In addition, Kowalczyk et al. point out that intragenic enhancers can act as alternative promoters^36^. Our DICs did not overlap with any known alternative promoters. Even though we know little about the molecular mechanism, our results strongly suggest a previously undescribed function of cohesin in intragenic regions with respect to gene expression regulation.

In summary, large-scale multi-omics enabled us to identify a cluster of cohesin DICs in MCF-7 and other cell types. Some tissue-specific DICs (LC-DICs) were related to enhancers and Pol2 pausing/release, whereas others (HC-DICs) contributed to chromatin architecture and might attenuate transcription. Our integrated analysis and machine learning approaches indicated distinct characteristics that distinguish DICs from other cohesin binding sites. Based on these genomic, epigenomic and transcriptomic characteristics, we can infer that DICs have distinct functions as compared with other cohesin sites.

## Methods

### Cell culture and treatment

RPE cells^37^ and MCF-7 cells (JCRB Cell Bank) were cultured in DMEM containing 10% FBS and 1% penicillin/streptomycin. Before subsequent treatments, RPE cells were cultured in serum-free medium for 48 hours and then were incubated in DMEM containing 10% FBS for 30 min. MCF-7 cells were maintained in phenol red–free medium containing charcoal-dextran–stripped FBS (Life Technologies) at 70–80% confluency for 2 days before treatment with 50 nM E2 (SIGMA) for the indicated times. Rad21 siRNA oligos^4^ were transfected using Lipofectamine RNAiMax (Life Technologies) according to the manufacturer’s instructions at 40 hours before treatment with E2. DRB (TCI) was added at 1.5 hours before treatment with E2. The effect of cohesin (Rad21)-deficiency were verified by western blot as shown in Supplementary Fig. 6e.

### ChIP and antibodies

Cells were fixed in medium or phosphate buffered saline with 1% formaldehyde at room temperature for 10 min. ChIP experiments were performed as described^38^. ChIP-seq libraries were prepared using NEBNext ChIP-seq Library Prep Master Mix Set for Illumina (New England BioLabs). Rabbit polyclonal antibody for Rad21 has been described in^39^. Antibody for Pol2 (#14958) was from Cell Signaling Technology. Antibodies for Mau2 (ab46906) and SA1 (ab4457) were from Abcam. Antibodies for TAF (A303-505A) and AFF4 (A302-538A) were from Bethyl Laboratory. CTCF (07-729) antibody was from Merck Millipore. Antibodies for Pol2ser2 and H3K27ac were kindly provided by Dr. H Kimura (TITech). Antibodies for P300 (sc-585) and Med1 (sc-5334) were from Santa Cruz Biotechnology. Antibody for CBP (606402) was from BioLegend.

### ChIP-seq analysis

After quality check by FastQC and SSP^40^, ChIP-seq reads were aligned to the human reference genome (hg38) using Bowtie^41^ version 1.2.2 with “-n2 -m1” parameters, by which we considered only uniquely mapped reads and allowed two mismatches in the first 28 bases per read. Peak calling was performed using MACS2^42^ version 2.2.6 with default settings. We used DROMPA^43^ version 3.7.2 to conduct statistical analysis and visualization. For visualization of ChIP-seq binding to particular chromatin regions, reads were normalized relative to total read number, and gene annotation was obtained from NCBI reference sequences (RefSeq; hg38). Read profiles around the sites of interest were plotted with the PROFILE mode of DROMPA, whereas the heatmap of target sites (2.5 kb around the peak summit) was plotted using HEATMAP mode. Genomic distribution in Fig. 2e was plotted by ChIPseeker^44^. Downstream analysis, such as peak overlap, was performed by Bedtools^45^ version 2.29.2 and Samtools^46^ version 1.9. Sources for all ChIP-seq data and other next-generation sequencing data (including our data and public data) are listed in Supplementary Tables S2-3.

### Hi-C analysis

All in-situ Hi-C data (control or E2-treated MCF-7 cells with two replicates) were aligned to the hg38 human reference genome. Further analysis was carried out mainly by Juicer^22^ version 1.11.04. All contact matrices were normalized by the KR method in Juicer. Chromatin loops were annotated using the HiCCUPS algorithm with default parameters^22^. The loop regions we used were merged from the results of 5-kb,10-kb and 25-kb resolutions. Aggregate peak analysis (APA) was performed using the ‘apa’ mode of Juicer (5-kb resolution), to measure the enrichment of the Hi-C signal around a set of peaks. The visualization of the contact matrix on the *MREG* locus was accomplished by Matplolib. After correction and normalization, comparable contact matrices were plotted at a 5-kb resolution. We merged two adjacent bins for smoothing. Other Hi-C analyses were performed using HOMER^10^. We made the Tag directory with the “GATC” restriction site sequence. Chromatin compaction scores ΔDLR and ΔICF were calculated for each 5-kb region across the genome (-res 5000) from a 15-kb window size (-window 15000). Other metrics including PC1, insulation score and TAD boundaries were obtained using HOMER with default parameters. We used the WashU epigenome browser^47^ to visualize Supplementary Fig. 3b.

### ChIA-PET analysis

RNA polymerase II–bound chromatin interactions in MCF-7 cells were extracted from ChIA-PET data (GSE33664). All fastq files were applied to the published pipeline Mango^48^ with default parameters, based on the hg38 reference genome. ChIA-PET interactions were visualized by DROMPA with the parameter ‘-inter’.

### RNA-seq and GRO-seq analysis

Using HISAT2^49^ version 2.2.0, we aligned paired-end RNA-seq reads to the index established from the hg38 reference genome. The output SAM files were converted to BAM files by Samtools. Htseq^50^ version 0.11.3 was then used with default parameters to generate a count table, which describes the number of reads on each gene. We used a GTF file (GRCh38.p12) from GENCODE for gene annotation. Subsequent differential expression analysis was achieved using DESeq2^51^, with its internal normalization. For GRO-seq, alignment was carried out using Bowtie with “-n2 -m1” parameters. The output was preprocessed and visualized using DROMPA.

### Data collection and machine learning

All datasets used in machine learning are listed in Table S5. Apart from our data, public omics data in wild-type MCF-7 cells were downloaded mainly from the GEO database, ENA database, ENCODE project, FANTOM5 project, UCSC genome browser and GWAS Catalog database. These data were then overlapped with all 184,140 cohesin peaks. As a result, we obtained 15 continuous features and 160 binary features, the latter of which indicated whether a kind of data was co-localized (1) or not co-localized (0) at a cohesin site. After normalization of continuous features, the big matrix, which consisted of 184,140 rows (cohesin sites) and 175 features, was imported into for different analyses. The features correlation heatmap for all cohesin sites and DICs was made with the R package *corrplot*. We used scikit-learn version 0.22.1 to perform machine learning. Overall, the parameters used in scikit-learn were optimized by grid search with 5-fold cross-validation. For unsupervised learning (k-means), all 175 features were used to fit models. For supervised learning (logistic regression, support vector machine, random forest), we omitted Mvalue, cohesin location and CTCF signal information and then used the remaining 168 features as independent variables *X*_*i*_ = (*X*_*i*1_, *X*_*i*2_, …, *X*_*ij*_), for *i* = 1, 2, …, 184140 and *j* = 1, 2, …, 168. Based on whether they were DICs or not, we labeled each cohesin site as 1 or 0 and then used it as a dependent variable *Y*_*i*_ ∈ {0, 1}. The conditional probability of logistic regression was calculated as follows:

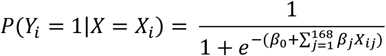

where β_*j*_ is the regression coefficient of each feature. We used training data to do model fitting and used test data to validate model performance. To apply the MCF-7–derived ML model to RPE cells, we used 25 features that were available in both MCF-7 and RPE cells. We used logistic regression with L1 penalty to decide whether each intragenic cohesin site had decreased binding (1) or not (0). Then the MCF-7–fitted model was applied to the RPE features to predict DICs.

We also applied penalized regression followed by univariate linear regression as described^52^, to reveal which features contributed to negative or positive Mvalue (log ratio of cohesin peak intensity between E2 and control) in intragenic cohesin (26066 sites). We used 169 features (of the 175 features, 6 were excluded: 5 features related to cohesin position and the Mvalue feature) as independent variables *X*_*i*_ = (*X*_*i*1_, *X*_*i*2_, …, *X*_*ij*_), for *i* = 1, 2, …, 26066 and *j* = 1, 2, …, 169, whereas the Mvalue was the dependent variable *Y*_*i*_. Instead of using the ordinary least squares approach, we used the elastic net loss function:

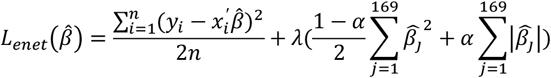

to the linear model 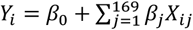, where *n* = 26066 and 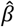 was the estimation of *β*. λ was chosen by cross-validation, and *α* = 0.5 was used to consider both the L1 and L2 penalty. Feature selection with such regularization was useful for filtering out non-significant or redundant features. The remaining features were applied to univariate linear regression *Y* = *a* + *bX* to calculate the regression coefficient (Supplementary Figs. 8f–h).

### Extraction of DICs

Quantitative comparison of Rad21 binding events was performed using MAnorm^19^ version 1.3.0 with default parameters. This results in the normalized Mvalue, a quantitative measure of differential binding for all cohesin sites. To acquire more convincing sites, we combined our results with high-quality ChIP-seq data from E-TABM-828^5^. Criteria for selecting decreased peaks were peak width < 3 kb and Mvalue < −0.5. Next, we used RefSeq as the reference genome to obtain intragenic regions. As described in Supplementary Fig. 1e, we excluded 10kb flanking regions around TSS and TES. Only large genes (gene length > 20 kb) were considered. E2-responsive genes were defined as genes with an increased Pol2ser2 ChIP-seq signal (ratio > 1.2) in the presence of E2 relative to control and that were validated by RNA-seq data. Decreased peaks at intragenic region of 499 E2-responsive gene were defined as DICs. Next, to quantify CTCF read density on DICs, we used MULTICI options in DROMPA software. Peaks with extremely low Rad21 signals were omitted. Finally, we separated the DICs into 141 high-CTCF DICs (HC-DICs) and 417 low-CTCF DICs (LC-DICs).

### Motif analysis

Motifs were analyzed using HOMER. Briefly, peak files in standard bed format were converted to HOMER peak files, and then the command *findMotifsGenome*.*pl* was used to discover the motif. The results included known motifs as well as *de novo* discovered motifs. The size of the region used for motif finding was set to 200 bp. The top 10 motifs with the lowest *q*-values (Benjamini-Hochberg) are shown.

### Software environment

All analyses were based on Ubuntu 18.04.4 with Python 3.6.9 and R 3.6.3. Data were processed using R base package or Numpy (v 1.17.2) as well as Pandas (v 0.25.1) in Python. Figures were drawn with DROMPA (v 3.7.2), Matplotlib (v 3.1.1), ggplot2 and R base plotting.

## Abbreviations

ChIP-seq: chromatin immunoprecipitation followed by sequencing
CNCs: cohesin-non-CTCF sites
RNA-seq: RNA sequencing
Hi-C: high-throughput chromosome conformation capture
ChIA-PET: chromatin interaction analysis by paired-end tag sequencing
DIC: decreased intragenic cohesin
LC-DIC: low-CTCF decreased intragenic cohesin
HC-DIC: high-CTCF decreased intragenic cohesin
Pol2: RNA polymerase II
Pol2ser2: RNA pol II CTD serine-2 phosphorylation
TSS: transcription start site
TES: transcription end site
TAD: topologically associating domain

## Author contributions

J.W. performed all bioinformatics analysis. R.N. conceived this project. J.W. and R.N. drafted the manuscript. M.B. prepared ChIP-seq and RNA-seq samples. K.S. supervised the sample preparation and sequencing, suggested ways to improve the analysis, and improved the manuscript.

## Data availability

The raw sequencing data and processed files are available at the GEO

